# Predicting uptake and elimination kinetics of chemicals in invertebrates: a technical note on residual variance modeling

**DOI:** 10.1101/2024.10.14.617565

**Authors:** Henk J. van Lingen, Edoardo Saccenti, Maria Suarez-Diez, Marta Baccaro, Nico W. van den Brink

## Abstract

Toxicokinetic models for predicting contents of nanomaterials and other toxic chemicals are often fitted without evaluation of the residual variance structure. The aim of the present study was to evaluate various residual variance structures, assuming either homoscedasticity or heteroscedasticity, when fitting non-linear toxicokinetic one-compartment models for predicting uptake, bioaccumulation and elimination of chemicals in invertebrate organisms. Data describing the exposure of several aquatic and terrestrial invertebrates to specific metal nanomaterials and other chemicals were available from real experiments for evaluating the residual variance functions for toxicokinetic models. As proof of concept, datasets of truly homoscedastic and heteroscedastic nature were simulated. Depending the dataset, applying models with different residuals variance assumption largely affected the residual plots and the error margins of parameters or the predicted content of a chemical. Consequently, selecting the most accurate residual variance functions for toxicokinetic modeling, either homoscedastic or heteroscedastic, improves the prediction of chemical contents in invertebrate organisms and the estimation of the associated uptake and elimination rates.

**Highlights:**

- Residual plots indicate if an accurate model was fitted to the toxicological data
- Choice of residual variance function affects the error margins of predicted chemicals
- Selecting proper residual variance models may prevent false positives and negatives

## 1. Introduction

Toxicokinetic modeling is a widely applied quantitative technique in the field of ecotoxicology. Fitting these models enables the estimation of uptake and elimination rates along with bioaccumulation of nanomaterials and other potentially toxic chemicals in environmental organisms, which is an important aspect of environmental risk assessment (e.g. Ref. 1). Accumulation of chemicals depends on the concentration in exposure media and physiological traits of the organism, which may be reflected by the estimated uptake and elimination rates (2). Modeling these toxicokinetic processes dynamically is often referred to as physiology-based pharmocokinetic modeling or biodynamic modeling. Fitting non-linear one-compartment toxicokinetic models is a relatively straightforward approach that considers organisms as a single compartment, describing only uptake and elimination (3). Sometimes, two or more compartments are considered to account for storage of chemicals, for multiple organs, for metabolizing to other metabolites, or for the internal distribution of the chemical within an organism (4; 5). However, a multi-compartment approach requires a more complex model and accurate and precise data for multiple compartments (3). Therefore, the one-compartment model appears the most commonly applied toxicokinetic approach dealing with various terrestial invertebrate species (e.g. Ref. 6; 7; 8) and aquatic species (e.g. Ref. 9; 10; 11) exposed to different types of nanomaterials and other toxic chemicals.

Although one-compartment non-linear models are commonly applied, estimating model parameters and their uncertainty can be challenging, as these estimates depend on the residual variance structure of the model (12). Ordinary-least-squares optimization assumes homoscedasticity (i.e. equal variance), which results in biased estimates of parameter variances when applied to heteroscedastic data, thereby invalidating significance testing of model parameters and variables (13). When heteroscedasticity occurs in biological assay data, observations with the greatest variance contribute proportionally more to the sum of squares, biasing the fit towards those points of greatest variance (14). Furthermore, recent studies have emphasized the importance of uncertainty estimates of toxicokinetic models for environmental risk assessment (15; 16; 17), leading to the development of the rbioacc R-package for analyzing toxicokinetic data (18). Despite this commendable effort, uncertainty estimation was not assessed in relation to heteroscedasticity. Data transformation could improve the residual variance distribution (19; 20; 21), but may not always be the most accurate approach (22). Residual variance modeling may then be a viable alternative (20; 23; 24).

The aim of the present study is to evaluate various residual variance functions when fitting non linear toxicokinetic models for predicting uptake, bioaccumulation and elimination of nanomaterials and other chemicals in invertebrate organisms.

## 2. Materials and Methods

### 2.1. Non-linear toxicokinetic models and residual variance functions

A one-compartment toxicokinetic model (2) was taken for evaluating various residual variance assumptions. This model describes an uptake and elimination phase, with the uptake phase given by Eq. 1:

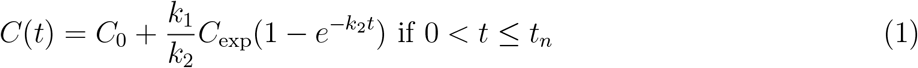

and the elimination phase given by Eq. 2:

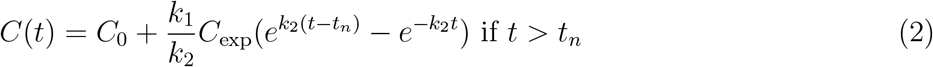

where *C*(*t*) represents instantaneous content of the chemical at time *t* in mg/g or *µ*g/g animal tissue, *C*_0_ represents the basal internal chemical content at *t* = 0 in *µ*g/g animal tissue (mg or ng rather than *µ*g may apply for certain systems), *k*_1_ represents the uptake rate in g soil/g animal tissue per time unit (e.g. hour, day) or L medium solution/g animal tissue per time unit, *k*_2_ represents the elimination rate per time unit, the 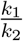 fraction is considered the kinetic bioaccumulation factor (BAF; g/g or L/g) and quantifies the ratio of the chemical contents in the animal divided by the exposure medium, *C*_exp_ represents the chemical concentration in the exposure medium in mg/kg, mg/L or *µ*/kg, et cetera, and *t*_*n*_ represents the time (day or hour) of transfer of the organisms from exposed to clean media. This one-compartment toxicokinetic model was fitted as a non-linear regression model according to (Eq. 3):

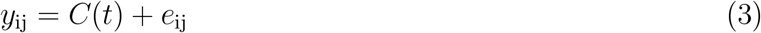

where *y*_*ij*_ represents the observed chemical content *j* at time *i* with *e*_*ij*_ being the corresponding residuals. These residuals are assumed to be homoscedastic (i.e. equal variance) or heteroscedastic (i.e. non-equal variance) and may be described by various residual variance function (Ref. (24)). Hence, if homoscedasticity is assumed, residual variances for the considered non-linear model fits are described according to (Eq. 4):

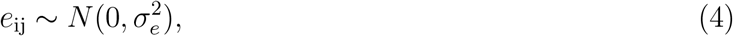

where 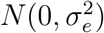 represents a normal distribution with zero mean and error variance 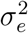 However, for heteroscedastic residual assumptions, the residual variance may increase along increasing *y*_*ij*_ using fixed variance according to (Eq. 5):

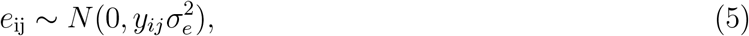

using exponential variance according to (Eq. 6):

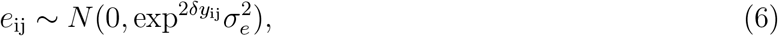

or using power variance according to (Eq. 7):

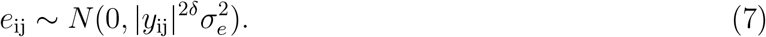

In addition, the one-compartment toxicokinetic model was fitted as a non-linear regression model as described by (Eq. 8):

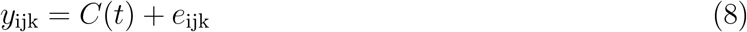

where *y*_ijk_ represents the chemical content *j* at time *i* in phase *k*, with *e*_ijk_ being the corresponding residuals, while assuming different residual variances for the uptake and elimination phases using stratified variance according to (Eq. 9):

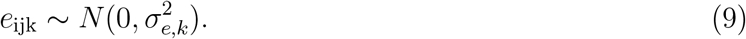

Here, *k* residual variance parameters are estimated separately for the uptake and elimination phases.

### 2.2. Experiments and datasets

#### 2.2.1. In-house data for earthworms exposed to ZnO:Mn and MnCl_2_

An *in vivo* toxicokinetic experiment was performed with earthworms, ordered as *Eisenia fetida* specimens (LASEBO, Nijkerk, The Netherlands), in glass jars containing LUFA 2.2 reference soil (LUFA Speyer, Germany). Based on the information provided by the supplier, the test soil had a pH of 5.6 *±* 0.29 (mean *±* SD) that was determined using 0.01 M CaCl_2_, an organic carbon content of 1.82 *±* 0.48% (w/w), a measured cation exchange capacity of 9.54 *±* 1.36 meq/100 g, and a water holding capacity of 48.9 *±* 5.6 %. The soil was air-dried and *<*2 mm sieved.

The earthworms were exposed to wet soil (50% WHC) homogeneously contaminated with a ZnO:Mn multicomponent nanomaterial and to MnCl_2_. The nominal exposure content of ZnO:Mn was set at 850 mg/kg soil, resulting in a Zn content equal to 650 mg Zn/kg soil. This content was chosen to ensure quantifiable zinc above the natural background content in earthworms. The nominal exposure content of Mn equal to 45 mg/kg soil corresponded to the Zn/Mn ratio used for the production of the ZnO:Mn nanomaterial (95% ZnO and 5% Mn). Furthermore, to match the manganese content in ZnO:Mn treated soil, the manganese content for MnCl_2_ treated soil was also set at 45 mg Mn/kg soil.

Throughout the entire incubation period, jars containing soil and clitellated earthworms were stored in a climate-controlled cabinet (Weiss Technik, Germany) at 20°C, 75% air humidity and a light intensity equal to 30 *µ*mol/m^2^/s. Lights were switched on for 24 hours per day to stimulate the earthworms to stay in the soil and to avoid any escape from the jars. Worms were fed with horse manure from an organic farm (Bennekom, The Netherlands) with known absence of pharmaceutical use. Whole worm samples were collected at days 1, 2, 7, and 14 during the uptake phase. After 14 days of exposure, the worms of the remaining jars were transferred to jars with clean LUFA 2.2 reference soil (n = 4 for each time point) for the elimination phase. Additional sample collection took place on days 15, 16, 21 and 28 days from the start of the exposure. At each sampling time earthworms were depurated from their gut content for 24 hours and snap frozen using liquid nitrogen.

Following homogenization and freeze-drying, similarly to (Ref. 25), Zn and Mn extraction from earthworm tissue was performed using Teflon vessels and 7 mL of reverse aqua regia (3:1 ratio of HCl:HNO_3_; Merck, Darmstadt) on hot plates in an open system. After diluting properly, the digested samples were analyzed by inductively coupled plasma-mass spectrometry (ICP-MS Nexion 350D, Perkin-Elmer Inc., Waltham, MA). The calibration curve was prepared from solutions of Zn^2+^ and Mn^2+^ (standard stock solution 1000 mg/L of ionic metal ion, Merck, Darmstadt) in a matching matrix.

#### 2.2.2. Acquired data for amphipod crustacean exposed to propanolol and springtails exposed to copper

Literature data for *Gammarus pulex* (an amphipod crustacean) exposed to propanolol for 48 hours and then transferred to a clean environment for another 48 hours ((see Ref. 26, for experimental details)) were extracted using https://apps.automeris.io/wpd4/. Furthermore, data was extracted for *Folsomia candida* (a springtail) exposed to Cu for 14 days and then transferred to clean environment for another 14 days ((see Ref. 27, for experimental details)). Briefly, *G. pulex* and *F. candida* were exposed to concentrations of propanolol and copper equal to 0.92 *µ*g/L water and 100 *µ*g/g soil, respectively. From the start of the exposure, sampling took place at 2, 5, 18, 24, 48, 50, 53, 66, 72 and 96 hours for *G. pulex* and at 0.25, 1, 4, 7, 10, 14.25, 15, 16, 18, 21, 24 and 28 days for *F. candida*. Additionally, data was obtained from experiments in which *F. candida* were exposed to contaminated soil that contained 25 *µ*g Cu/g soil (27) and 20 mmol Cd/g soil with pH values of 4.5, 5,5 and 6.5, respectively (6). Sampling was also performed at 0.25, 1, 4, 7, 10, 14.25, 15, 16, 18, 21, 24 and 28 days from the start of the exposure and the transfer to clean soil was on day 14.

#### 2.2.3. Simulated data for invertebrate species exposed to chemicals

Toxicokinetic data was simulated to describe any chemical content in any organism in time along uptake and elimination phases. For this data simulation, time ran over 28 units of time (e.g. days, hours, seconds) with *t*_*n*_ indicating the transfer from exposure to a clean environment at time unit 14. Parameters in the one-compartment model as represented in Eqns. 1-2 for the uptake and elimination phases were set as: *C*_0_ = 5 mg/g, *k*_1_ = 10 mg/g per time unit and *k*_2_ = 1 per time unit, along with the chemical content in the exposure medium *C*_exp_ = 1 mg/g. Setting these parameters and variable enabled the computation of error-free chemical contents that reached a plateau obviously above basal level along the uptake phase and declined to basal level again long the elimination phases. A simulation was then performed to obtain data with homoscedastic residual variance using *e*_ij_ *∼* uniform(*™*3, 3) and to obtain data with heteroscedastic residual error variance using *e*_ij_ *∼ N* (0, *y*_ij_ *·*0.2) and had a sample size *n* = 4. Simulated data were uncorrelated, meaning no repeated measurements in time were taken from the same experiment unit (e.g. jar with soil, tank with water, etc). After generating datasets with homoscedastic and heteroscedastic residual variances for all 28 time points, reduced homoscedastic and heteroscedastic datasets were obtained by only retaining data for time units 1, 2, 7, 14, 15, 16, 21 and 28.

### 2.3. Model fitting and Bayesian implementation

The homoscedastic model and the various heteroscedastic models were fitted within a Bayesian framework. For these Bayesian models, informative priors were specified as *C*_0_, *k*_1_, *k*_2_ *∼ N* (*µ, σ*) with *µ* and *σ* being the parameter means and standard errors taken from frequentist (i.e. non-Bayesian) homoscedastic model fits. Additional priors were *σ*_*e*_, *σ*_*e,k*_ *∼* half-Cauchy(0,5) and *δ ∼ N* (*µ* = 0, *σ* = 1000). The choice of prior distributions for *σ* and *δ* aimed to construct minimally informative priors. For each model that was fitted, 4 chains were run for 53 *×* 10^3^ iterations with the first 3 *×* 10^3^ taken as burn-in period. The chains were then thinned by a factor of 25, after which convergence was assessed by visually inspecting the chain traces of the various model parameters.

Following the fitting of the various models, posterior predictive distributions (PPD; (e.g. Ref. 28)) were simulated. For the linear, exponential and power residual variance models, which assume the error variance is dependent on the magnitude of the chemical content, the error variance was computed using the mean predicted values of the chemical rather than the observed values. Moreover, the goodness-of-fit relative to the number of estimated parameters per model was computed using the leave-one-out cross-validation information criterion (LOOIC; (29)).

Model simulations were run using the `nls()` function in R (version 4.2.2; Ref. (30)), interfaced using Rstudio (version 2022.07.0; Ref. (31)) and through the `rstan` (Stan Development Team, 2016) and loo packages (32). All R and STAN code for fitting the various models is available at https://github.com/linge006/FittingToxicokineticModels. Additionally, for those who prefer not to apply Bayesian statistics, R code is available for frequentist fitting of the various models using the gnls() function from the nlme package(33).

## 3. Results

### 3.1. Earthworms exposed to ZnO:Mn and MnCl_2_

Earthworms were exposed to soils containing the ZnO:Mn nanomaterial and MnCl_2_ that was used as control for 14 days (uptake phase), followed by 14 days of exposure to clean soil (elimination phase). Measured zinc and manganese contents in earthworm tissue were used to fit a one-compartment toxicokinetic model along uptake and elimination phases (Eqns. 1 and 2, respectively) with the residual variance models indicated in Eq. 4 (homoscedastic model) and Eqns. 5 to 7 and 9 (heteroscedastic models). Most data points for the zinc content in earthworms exposed to ZnO:Mn were within the bounds of the posterior predictive distribution (PPD) of the homoscedastic model throughout the uptake phase (Fig. 1; see Table 1 for the posterior means of the parameters along with the log likelihood and LOOIC, the latter of which contributes to the selection of the best model residual variance function). The PPD provides an estimate of the distribution of future data observations given both the observed data and the model parameters, and is commonly used to check the consistence of the model with the data (28). However, the data points throughout the elimination phase show less variation than the bounds of the PPD of the homoscedastic model suggest. Therefore, the data does not appear homoscedastic and may require a heteroscedastic model. Hence, for the stratified heteroscedastic model, which indicated greater error variance throughout the uptake phase than the elimination phase, the ranges of the data points align with the bounds of the PPD for both phases. This suggests the stratified variance model best fits these data that have a heteroscedastic nature, which was also confirmed by the logLik and LOOIC scores.

**Table 1:** Estimated parameters (mean *±* standard deviation) for the fitted toxicokinetic homoscedastic and selected heteroscedastic models for predicting chemical contents along with log likelihood (logLik; quantifies the goodness of fit for which a higher score indicates better fit) and leave-one-out information criterion (LOOIC; quantifies the goodness of fit relative to the number of model parameters for which a lower score indicates a better fit). *C*_0_ represents the basal internal chemical content in *µ*g/g (mg or ng rather than *µ*g may apply for certain systems), *k*_1_ represents the uptake rate in g soil/g animal per time unit (e.g. hour, day) or L medium solution/g animal per time unit, *k*_2_ represents the elimination rate per time unit (/h or /d), BAF represents the bioaccumulation factor in g/g or L/g. Model output is shown for the various in-house, acquired and full and reduced simulated datasets

| Model fit | $C_0$ | $k_1$ | $k_2$ | BAF | $\sigma_e$ or $\sigma_{e,\text{uptake}}$ | $\sigma_{e,\text{elimin}}$ or $\delta$ | logLik | LOOIC |
| --- | --- | --- | --- | --- | --- | --- | --- | --- |
| *E. fetida* exposed to ZnO:Mn (in-house data) |  |  |  |  |  |  |  |  |
| Homoscedastic Zn | 69.2±8.4 | 0.711±0.193 | 3.19±0.91 | 0.224±0.022 | 44.2±5.7 | | -169 | 339 |
| Stratified Zn | 70.7±1.8 | 0.899±0.171 | 4.28±0.73 | 0.211±0.024 | 63.2±11.9 | 7.0±1.4 | -145 | 291 |
| Fixed Zn | 69.4±6.2 | 0.692±0.180 | 3.57±0.94 | 0.195±0.020 | 3.32±0.44 | | -163 | 326 |
| Exponential Zn | 69.8±2.2 | 0.456±0.127 | 4.48±1.09 | 0.102±0.017 | 3.24±1.18 | 0.014±0.002 | -152 | 304 |
| Power Zn | 68.9±2.7 | 0.547±0.153 | 4.27±1.04 | 0.129±0.024 | 0.04±0.21 | 1.66±0.31 | -153 | 305 |
|  |  |  |  |  |  |  |  |  |
| Homoscedastic Mn | 40.5±2.3 | 0.052±0.030 | 1.64±0.81 | 0.036±0.020 | 14.2±1.8 | | -132 | 265 |
| Stratified Mn | 40.6±2.0 | 0.051±0.031 | 1.67±0.80 | 0.034±0.020 | 18.0±3.4 | 9.6±1.8 | -130 | 260 |
| Fixed Mn | 37.1±2.6 | 0.026±0.020 | 1.36±0.65 | 0.021±0.017 | 2.6±0.3 | | -137 | 274 |
| Exponential Mn | 39.1±2.8 | 0.037±0.025 | 1.30±0.60 | 0.032±0.021 | 11.7±2.4 | 0.0055±0.0045 | -133 | 266 |
| Power Mn | 39.5±2.7 | 0.040±0.025 | 1.27±0.60 | 0.035±0.021 | 9.9±3.2 | 0.12±0.10 | -134 | 267 |
|  |  |  |  |  |  |  |  |  |
| *E. fetida* exposed to MnCl<sub>2</sub> (in-house data) |  |  |  |  |  |  |  |  |
| Homoscedastic Mn | 33.2±1.7 | 0.115±0.041 | 2.31±0.84 | 0.052±0.014 | 9.0±1.2 | | -117 | 234 |
| Stratified Mn | 33.3±1.7 | 0.125±0.050 | 2.55±1.04 | 0.051±0.015 | 10.2±1.9 | 8.2±1.5 | -118 | 235 |
| Fixed Mn | 32.3±1.6 | 0.121±0.049 | 2.76±1.08 | 0.046±0.014 | 1.5±0.2 | | -117 | 233 |
| Exponential Mn | 33.2±1.6 | 0.114±0.042 | 2.27±0.85 | 0.052±0.015 | 10.1±1.9 | 8.1±1.5 | -116 | 233 |
| Power Mn | 32.5±1.8 | 0.124±0.049 | 2.73±1.06 | 0.048±0.014 | 3.2±2.3 | 0.37±0.24 | -117 | 234 |
|  |  |  |  |  |  |  |  |  |
| *G. pulex* exposed to propranolol (acquired data) |  |  |  |  |  |  |  |  |
| Homoscedastic | 0.92±0.66 | 0.563±0.054 | 0.018±0.003 | 32.6±4.4 | 3.72±0.52 | | -84 | 168 |
| Stratified | 0.95±0.68 | 0.549±0.058 | 0.017±0.003 | 32.5±4.5 | 4.13±0.85 | 3.48±0.76 | -85 | 169 |
| Fixed | 0.34±0.27 | 0.526±0.047 | 0.016±0.003 | 32.9±4.1 | 0.99±0.14 | | -74 | 149 |
| Exponential | 0.53±0.32 | 0.461±0.043 | 0.015±0.002 | 30.5±3.4 | 0.76±0.25 | 0.11±0.02 | -70 | 140 |
| Power | 0.26±0.25 | 0.515±0.047 | 0.016±0.003 | 32.9±4.0 | 0.81±0.32 | 0.61±0.14 | -75 | 150 |
|  |  |  |  |  |  |  |  |  |
| *F. candida* exposed to Cu at 100 mmol/g of soil with pH = 5.5 (acquired data) |  |  |  |  |  |  |  |  |
| Homoscedastic | 59.1±7.6 | 0.100±0.038 | 0.25±0.10 | 0.43±0.18 | 30.1±4.6 | | -113 | 225 |
| Stratified | 59.8±7.7 | 0.102±0.039 | 0.26±0.10 | 0.42±0.16 | 33.9±9.3 | 28.6±5.6 | -114 | 227 |
| Fixed | 52.0±7.1 | 0.088±0.038 | 0.26±0.10 | 0.37±0.29 | 3.6±0.6 | | -112 | 225 |
| Exponential | 50.3±7.4 | 0.074±0.037 | 0.27±0.11 | 0.30±0.19 | 12.2±5.2 | 0.012±0.005 | -112 | 224 |
| Power | 53.2±8.2 | 0.090±0.038 | 0.26±0.10 | 0.41±0.21 | 7.5±6.4 | 0.38±0.17 | -113 | 225 |
|  |  |  |  |  |  |  |  |  |
| Homoscedastically simulated data |  |  |  |  |  |  |  |  |
| Homoscedastic (full data) | 1.92±0.11 | 5.01±0.45 | 0.97±0.09 | 5.17±0.21 | 1.19±0.08 | | -185 | 370 |
| Stratified (full data) | 1.92±0.11 | 5.01±0.46 | 0.97±0.09 | 5.17±0.21 | 1.21±0.12 | 1.19±0.11 | -186 | 372 |
| Fixed (full data) | 0.96±0.09 | 5.55±0.51 | 0.94±0.09 | 5.90±0.34 | 0.93±0.07 | | -226 | 453 |
| Exponential (full data) | 1.89±0.11 | 5.02±0.46 | 0.98±0.09 | 5.14±0.21 | 1.09±0.10 | 0.021±0.016 | -186 | 371 |
| Power (full data) | 1.89±0.12 | 5.04±0.45 | 0.97±0.09 | 5.19±0.21 | 1.16±0.08 | 0.028±0.024 | -186 | 372 |
|  |  |  |  |  |  |  |  |  |
| Homoscedastic (red data) | 1.87±0.25 | 4.72±0.53 | 0.94±0.11 | 5.05±0.44 | 1.22±0.16 | | -52 | 104 |
| Stratified (red data) | 1.87±0.25 | 4.72±0.53 | 0.93±0.11 | 5.07±0.47 | 1.32±0.27 | 1.22±0.24 | -53 | 106 |
| Fixed (red data) | 1.15±0.20 | 5.07±0.58 | 0.92±0.10 | 5.52±0.56 | 0.79±0.11 | | -60 | 121 |
| Exponential (red data) | 1.78±0.24 | 4.72±0.53 | 0.95±0.11 | 5.00±0.47 | 1.00±0.20 | 0.051±0.039 | -53 | 105 |
| Power (red data) | 1.77±0.25 | 4.77±0.53 | 0.94±0.11 | 5.11±0.46 | 1.11±0.17 | 0.094±0.073 | -53 | 106 |
|  |  |  |  |  |  |  |  |  |
| Heteroscedastically simulated data |  |  |  |  |  |  |  |  |
| Homoscedastic (full data) | 2.07±0.09 | 6.67±0.72 | 1.44±0.16 | 4.63±0.16 | 0.96±0.06 | | -162 | 323 |
| Stratified (full data) | 2.04±0.06 | 5.89±0.62 | 1.25±0.13 | 4.72±0.19 | 1.32±0.13 | 0.47±0.05 | -137 | 274 |
| Fixed (full data) | 2.00±0.07 | 6.34±0.70 | 1.42±0.16 | 4.47±0.16 | 0.42±0.03 | | -139 | 278 |
| Exponential (full data) | 1.98±0.05 | 5.54±0.68 | 1.37±0.17 | 4.04±0.18 | 0.26±0.04 | 0.25±0.03 | -131 | 263 |
| Power (full data) | 1.92±0.05 | 5.97±0.70 | 1.38±0.16 | 4.34±0.18 | 0.21±0.04 | 0.97±0.12 | -131 | 262 |
|  |  |  |  |  |  |  |  |  |
| Homoscedastic (red data) | 2.05±0.22 | 6.67±0.68 | 1.27±0.14 | 5.28±0.40 | 1.08±0.14 | | -49 | 98 |
| Stratified (red data) | 2.04±0.18 | 6.56±0.71 | 1.24±0.14 | 5.29±0.42 | 1.42±0.28 | 0.73±0.15 | -47 | 93 |
| Fixed (red data) | 1.93±0.17 | 6.50±0.67 | 1.26±0.14 | 5.19±0.39 | 0.47±0.06 | | -44 | 88 |
| Exponential (red data) | 1.96±0.16 | 6.23±0.68 | 1.26±0.14 | 4.95±0.39 | 0.16±0.06 | 0.44±0.14 | -44 | 87 |
| Power (red data) | 1.87±0.17 | 6.44±0.69 | 1.24±0.14 | 5.19±0.40 | 0.39±0.15 | 0.68±0.23 | -44 | 88 |

**Figure 1:**
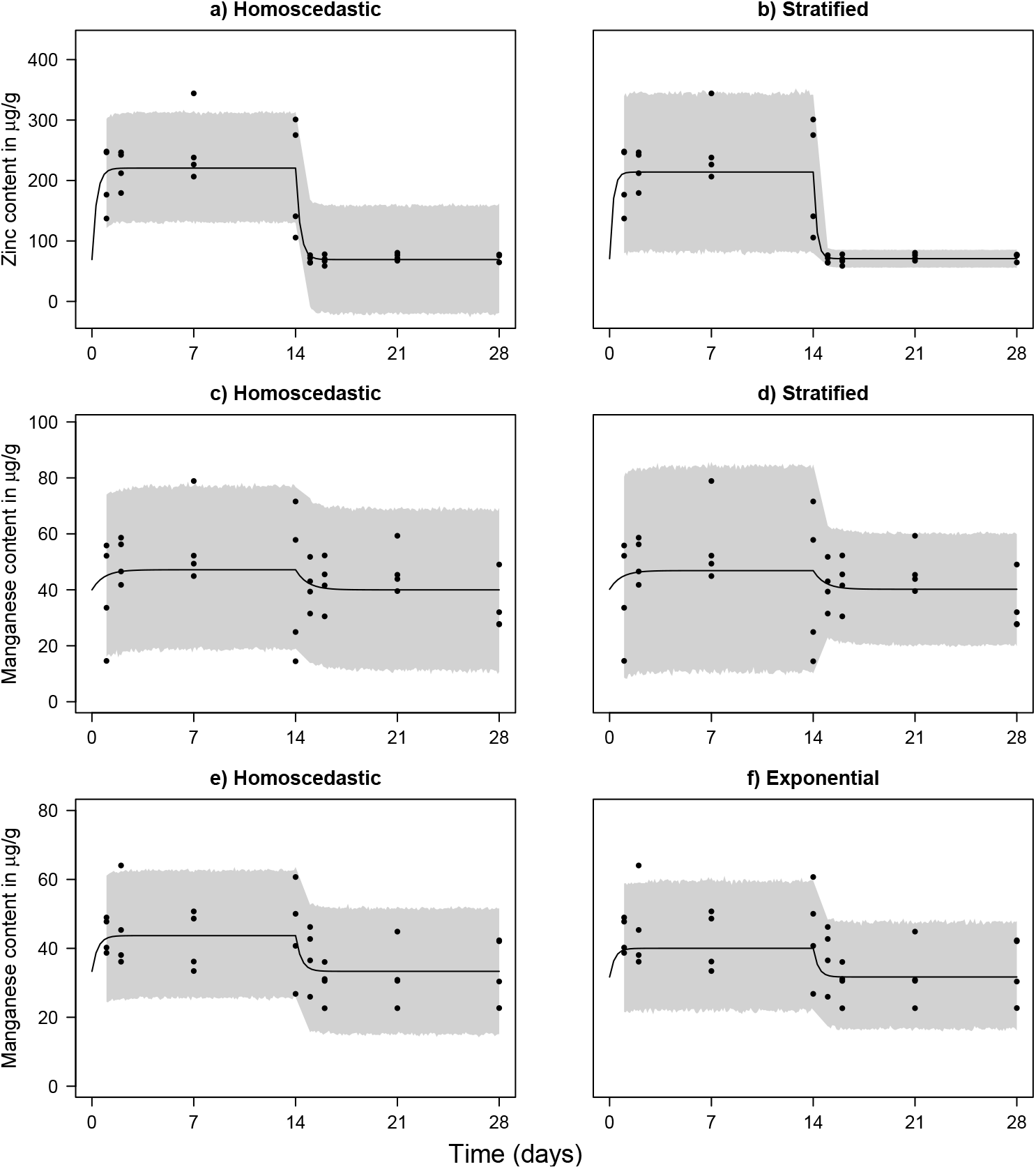
Fitted one-compartment toxicokinetic models assuming homoscedastic residual variance (panels a, c and e) and heteroscedastic residual variance (panels b, d and f) using the in-house *E. fetida* earthworm data. Data points represent the Zn and Mn contents in earthworms exposed to the ZnO:Mn nanomaterial (panels a and b, respectively), and the manganese contents in earthworms exposed to MnCl_2_ (panels e and f) throughout 28 days. The black line and the grey shaded area indicate the mean and 95% credible intervals of the posterior predictive distribution, respectively. Estimated model parameters and fit statistics are shown in Table 1

The data points for the manganese content in earthworms exposed to ZnO:Mn align better with the PPD bounds of the heteroscedastic model than the homoscedastic model. However, the lower PPD bound of the heteroscedastic model during the uptake phase is lower than during the elimination phase, which is not toxicologically feasible. The homoscedastic model fits the data points for the manganese content in earthworms exposed to MnCl_2_ relatively well, with only one point outside the PPD. The heteroscedastic model indicates two data points are outside the PPD, but aligns slightly better with the ranges of the data points during the elimination phase. Residual plots revealed the stratified residual variance model fit to the Zn contents in earthworms exposed to ZnO:Mn had a rather homogeneous pattern of variability for the standardized residuals, indicating this heteroscedatic model describes the within-group variance obviously better than the homoscedastic model (Fig. 2. However, the heteroscedastic model does not show a clear improvement over the homoscedastic model for manganese contents in the ZnO:Mn and MnCl_2_ exposed earthworms.

**Figure 2:**
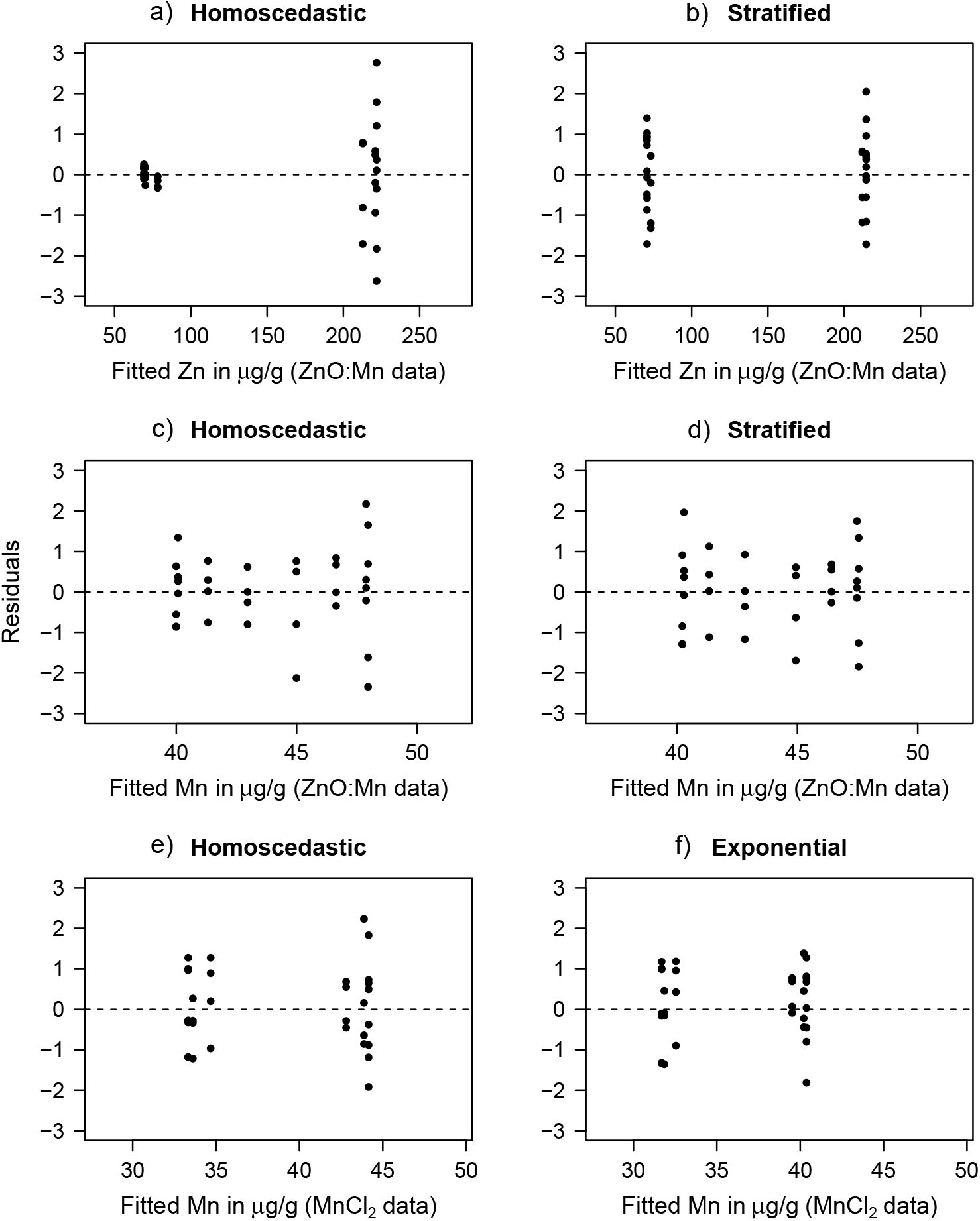
Residuals of the fitted one-compartment toxicokinetic models to the in-house data of Zn and Mn contents for *E. fetida* earthworms exposed to the ZnO:Mn nanomaterial (panels a-d, respectively), and the manganese contents in earthworms exposed to MnCl_2_ (panels e and f). Residuals are shown for the homoscedastic model fit (panels a, c and e) and the heteroscedastic model fits that used stratified, stratified and exponential residual variance functions, respectively (panels b, d and f).

### 3.2. Amphipod crustacean exposed to propanolol and springtails exposed to copper

The PPD of the homoscedastic model fit to the propanolol contents in *G. pulex* was wide enough to include nearly all data points, with only one data point at *t* = 14 beyond the upper bound (Fig. 3; see Table 1 for model parameter values). At *t* = 0 and *t* = 96, the PPD was wider than the ranges of the data. The exponential error variance model fit had a narrower PPD at the start and end of the time span of observations, aligning more closely with the data points at *t* = 0 and *t* = 96. However, the exponential error variance model also resulted in 2 data points, at 24 and 48 hours, respectively, outside the bounds of the PPD. Residual plots visualized the data points violated the equal variance assumption of the homoscedastic model, whereas the exponential residual variance model fitted the data well (Suppl. Fig. 1). One may note that performing 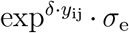 for the exponential model yields 3.96 for *y*_ij_ = 15 *µ*g/g, which is relatively similar to the values of *σ*_e_ obtained for the homoscedastic and stratified models. Evaluating the goodness-of-fit relative to the number of estimated parameters by LOOIC favored the exponential model by 28 score points. For the copper contents in *F. candida*, the homoscedastic model’s PPD bounds aligned well with the data ranges, though the lower bound was slightly below observed values during the elimination phase. The exponential residual variance model had a PPD for which a few points were above its upper bound, whereas the lower bound of the PPD was slightly below the various data points. Residual plots indicated the equal variance assumption of the homoscedastic model was not violated (Fig. A.5) and the LOOIC score indicated a minimal difference of only 2 points between the homoscedastic and exponential models. Additional examples of fitting one-compartment models using residual variance functions to data for *F. candida* exposed to copper and cadmium are provided in Fig. A.6 and Table A.2.

**Figure 3:**
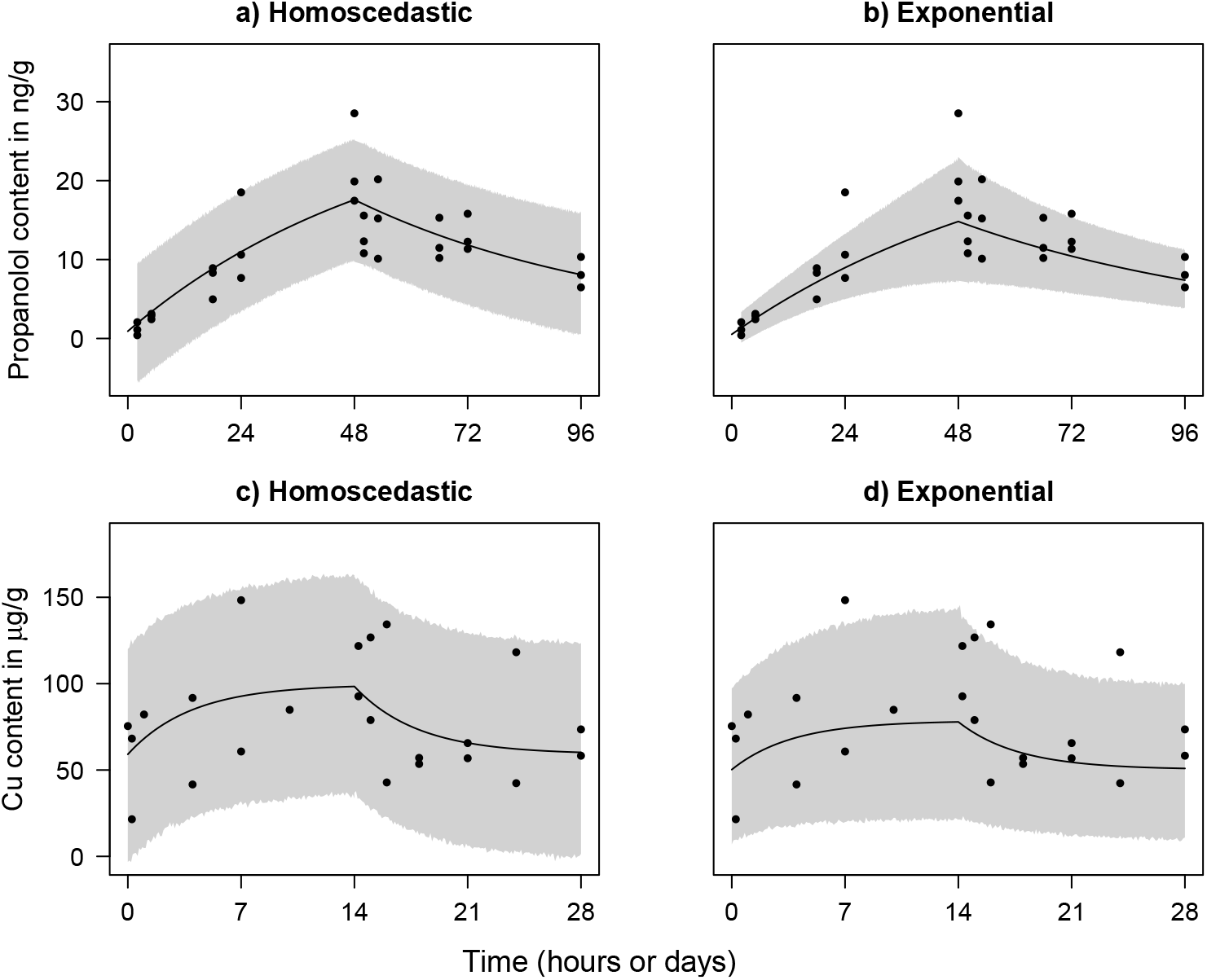
Fitted one-compartment toxicokinetic models assuming homoscedastic residual variance (panels a and c) and heteroscedastic residual variance (panels b and d) using the acquired data. Data points represent propanolol contents in ng/g wet weight of *G. pulex* throughout 96 hours (panels a and b) and copper contents in *µ*g/g dry weight of *F. candida* throughout 28 days (panels c and d). The black line and the grey shaded area indicate the mean and 95% credible intervals of the posterior predictive distribution, respectively. Estimated model parameters and fit statistics are shown in Table 1

### 3.3. Simulating invertebrate exposure to chemicals

The homoscedastic and exponential models had largely similar PPDs for the full homoscedastic dataset, with no data points outside the PPD (Fig. 4; see Table 1 for parameter values). Both models also fitted well to the reduced dataset, for which data points for days 1, 2, 7, 14, 15, 16, 21 and 28 were retained reflecting the earthworm data, although the PPD for the exponential model was slightly wider in the uptake phase and narrower in the elimination phase. LOOIC scores for the exponential model fits to the full and reduced data were marginally greater than for the homoscedastic model fits, indicating the homoscedastic model was preferred for the homoscedastically simulated data. Residual plots showed the homoscedastic residual variance model fit had a rather homogeneous pattern of variability for the standardized residuals, indicating a homoscedastic rather than a heteroscedatic model describes the within-group variance accurately (Fig. A.7).

**Figure 4:**
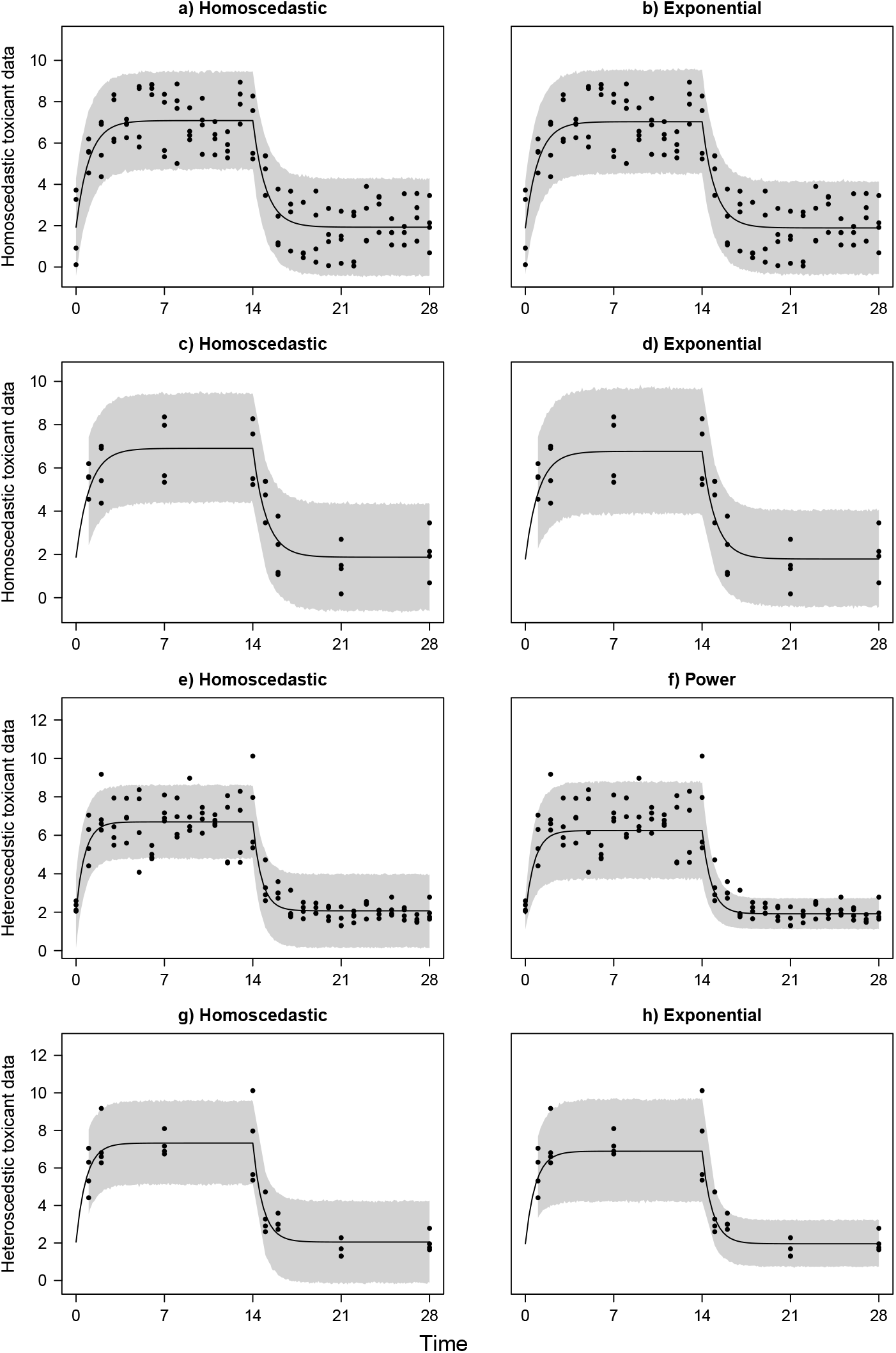
Fitted one-compartment toxicokinetic models assuming homoscedastic residual variance (panels a, c, e and g) and heteroscedastic residual variance (panels b, d, f and h) using simulated data. Data points represent contents of any chemical in any invertebrate throughout 28 time units and were simulated as homoscedastic (panels a-d) and heteroscedastic (panels e-h). Data points were retained for all 28 time units (full data; panels a, b, e and f) or only for time units 1, 2, 7, 14, 15, 16, 21 and 28 (reduced data; panels c, d, g and h). The black line and the grey shaded area indicate the mean and 95% credible intervals of the posterior predictive distribution, respectively. Estimated model parameters and fit statistics are shown in Table 1

For the full heteroscedastically simulated dataset, most data points were within the PPD bounds of the homoscedastic model, although some uptake phase points were outside the PPD, and the elimination phase data points ranged less than the PPD width. The power model had slighter better fit for which no uptake phase data points appeared below the lower bound of the PPD and the PPD aligning well with the range of the data points throughout the elimination phase. LOOIC scores confirmed the power model as the best fit to the data. The homoscedastic model fitted to the reduced heteroscedastically simulated data had one point outside the PPD at *t* = 14 and had a PPD that was wider than the range of the data points in the elimination phase. The exponential model fit to the reduced heteroscedastically simulated data had a PPD that was slightly wider in the uptake phase and narrower in the elimination phase. The latter PPD that aligned well with the data indicated the exponential model fitted the best to the data. For both the full and reduced heteroscedastically simulated datasets, LOOIC scores confirmed the best fits for the power and exponential models, respectively. Residual plots showed that the equal variance assumption of the homoscedastic model fits to the full and reduced heteroscedastically simulated data was violated (Fig. A.7). Fitting the power and exponential models to the full and reduced data, respectively, indicated relatively constant residual variance along the magnitude of the chemical contents.

## 4. Discussion

The present study uniquely explores the use of several residual variance functions for one-compartment non-linear toxicokinetic model fitting. Depending on the data, these residual variance functions affect the error margins of predicted values of the chemical content and the BAF. Various datasets, which were either of more homoscedastic or more heteroscedastic nature, were used to explore which residual variance function was the most accurate for each dataset. The various datasets available inhouse (earthworms) or acquired from the literature (Amphipod crustacean and springtail) described experimentally generated observations. For some of these datasets, the homoscedastic model fitted the data the best, whereas for other datasets, one of the heteroscedastic models fitted the data better. Furthermore, data of homoscedastic nature and data of heteroscedastic nature describing any chemical in any invertebrate were generated through *in silico* simulation. As proof of concept, the homoscedastic model fitted the best to the simulated truly homoscedastic data and a heteroscedastic model fitted the best to the simulated truly heteroscedastic data. These datasets represented terrestrial and aquatic invertebrates exposed to nanomaterials and other chemicals that are potentially toxic. All models applied in the present modeling effort were fitted using a Bayesian framework. Performing Bayesian simulation powerfully facilitated non-linear toxicokinetic model fitting and enabled the simulation of the PPD, which visualized the implications of the different residual variance functions. Furthermore, we provided R code for fitting the toxicokinetic one-compartment models using the various residual functions. These R codes are provided for both the context of Bayesian and frequentist (i.e. non-Bayesian) statistics.

The data compiled for this technical note reveal differences in chemical contents along the uptake and elimination phases. The variance for these observed contents may change throughout these phases, or may be specific for each of the two phases. Therefore, residual variance assumptions may be fulfilled differently within or throughout each phase, which must be considered when selecting the residual variance function for a toxicokinetic model fit. We have shown for the Zn data of the earthworms exposed to the ZnO:Mn nanomaterial that the residual variance function of the toxicokinetic models significantly affects the error margins of the posterior predicted zinc content. For example, testing whether the zinc content in earthworms is less than 100 *µ*g/g throughout the elimination phase would result in a false positive for the homoscedastic model, whereas the stratified model may yield a true negative. Accurate hypothesis testing may be relevant when using contents of toxic chemicals as biomarkers to assess whole-organism responses (34) predict declines in biomass and species abundance (35) and evaluate various other toxicological effects.

Moreover, for the other experimentally generated datasets, we demonstrated that the chosen residual function affects the error margin of the predicted chemical as visualized by means of the PPD. For the dataset that represented the zinc content in earthworms, error variance increased along increased values of the chemical, which was also shown for the dataset that represented the propanolol content in *G. pulex*, although to a lesser extent. In more general toxicokinetic modeling terms, one may verify if the variance increased at the occasion of a plateau chemical content throughout the uptake phase that is much greater than the basal content. Therefore, the model fitting to the data for *G. pulex* exposed to propranolol also indicates that hypothesis testing at baseline contents of the chemical could lead to false positive conclusions.

In general, standard errors of model parameters and variables from (non-linear) regression can be biased, implying that the associated 95% credible intervals (the Bayesian equivalent of frequentist confidence intervals) and *P* -values may not be reliable (36), potentially leading to false positive or false negative toxicological conclusions. It remains to be reviewed to what extent heteroscedasticity occurs in toxicology, as various studies (e.g. 6; 7; 8; 37) do not report any evaluation of variance equality. However, the present examples with earthworms and *G. pulex* demonstrate that residual variance structures have implications and should always be verified. Additional evidence supporting the importance of selecting accurate residual variance functions could be found outside the field of toxicology. For instance, inappropriate residual variance assumptions when predicting aortic valve area using body surface area increased the risk of misclassifying infants with abnormal aortic valve areas as normal, and misclassifying older children with normal aortic valve areas as abnormal (38).

Simulation of data contributed to further exploration of toxicokinetic modeling for datasets generated by experiments that include an uptake and elimination phase with variances in chemical contents that are either or not related to the magnitude of the chemical content. The homoscedastic model best fitted the simulated full and reduced homoscedastic datasets, while a heteroscedastic model best fitted the simulated full and reduced heteroscedastic datasets. This finding demonstrates that the optimal model alignment corresponds with the nature of the data, potentially applying to experimentally collected data as well. In the present study, we selected the best model, assuming either homoscedastic or heteroscedastic error variance, by computing the LOOIC score. Although this is a commonly accepted and applied model selection metric, similar to the Akaike and Bayesian Information Criteria in frequentist statistics (39; 40), and used in the field of toxicology (e.g. Ref. 41; 42), the model favored by the LOOIC may not always be completely toxicologically or biologically feasible. For example, the LOOIC favored the stratified residual variance model for the manganese content in earthworms exposed to the ZnO:Mn. However, this model showed a PPD with a lower bound that was lower in the uptake phase than in the elimination phase, which is not toxicologically feasible. Then, the homoscedastic model may be preferred over the stratified residual variance model, despite a 5-point higher LOOIC score. Finally, it is important to recognize that there is no clear consensus on the various information criteria and the penalty for the number of parameters for model selection purposes (43), which suggests that LOOIC should be used as a guideline rather than a strict rule.

Despite the various criteria and considerations for selecting the most accurate residual variance function for toxicokinetic model fitting, differences between the fits of residual variance models may be small or arbitrary. For instance, the datasets describing chemical contents in earthworms exposed to MnCl_2_ and *F. candida* exposed to copper showed relatively similar logLik and LOOIC values, and PPDs for different models. Such similar model fits may sometimes occur and complicate model selection. Nonetheless, we recommend keeping models as parsimonious as possible, starting with the assumption of homoscedasticity. If residual plots do not indicate any violations of model assumptions, not even by homoscedasticity evaluation such as the Breusch-Pagan test as used for epidemiological data analysis (44), it is reasonable to adhere to the applied residual variance assumptions and consider the model in question. If the residual plot of a heteroscedastic model does not show any patterns or anomalies, it might only be preferred over the homoscedastic model if the residuals exhibit fewer patterns or if logLik, LOOIC, and PPD show a clear improvement. Nevertheless, this consideration may still not suffice in every case, as each dataset is unique and may require a custom approach.

Although perfect datasets can be simulated, they are not commonly obtained on-site. A particular model may fit individual data points better, whereas another may fit most other data points more accurately. In such cases, individual points should not be decisive. Researchers must recognize that many datasets may contain one or more outliers or data point that contain certain patterns not perfectly aligning with any distribution, and a perfect model may not always exist. Despite this unpredictable nature or lack of consistency of data, researchers in toxicology should always strive to generate the highest quality datasets as possible. When performing toxicokinetic experiments that include uptake and elimination phases, we recommend that the experimental design includes sufficient sampling times and number of samples per sampling time. This may prevent one or two outliers or specific sampling times of a dataset being too influential.

In the present toxicokinetic modeling effort, only one-compartment models were considered, although multi-compartment models and models that account for a stored fraction of a chemical in an organism are often applied as well (2). These other two types of toxicokinetic models are also associated with residuals when fitted as non-linear regression models, and the variance of these residuals may be homoscedastic or heteroscedastic. Therefore, the residual variance functions explored in the present study could apply to toxicokinetic models other than one-compartment models. Depending on the toxicological problem and the type of data, transformations such as the Box-Cox transformation may be effective strategies for analyzing heteroscedastic data in the field of toxicology (21). Accounting for heteroscedasticity using residual variance functions leaves the toxicokinetic forcing functions unchanged and retains the toxicological meaning of the various model variables and parameters. Additionally, non-parametric error estimation (45) or heteroscedasticity-consistent standard error estimation for unknown functional forms of heteroscedasticity (36) may occasionally be applied for analyzing toxicological data. The present paper highlights tools for applying residual variance functions and shares considerations that may help selecting the most adequate tool. Researchers may then apply these tools given the specific nature of their toxicological data and be enabled to perform their hypothesis testing such that the occurrence of false positives and false negatives (type I and II errors) is minimized.

## 5. Conclusion

The present paper demonstrates the use of various residual variance functions when fitting toxicokinetic models and may be relevant for researchers working in other disciplines as well. A collection of datasets describing several invertebrates exposed to various metal nanomaterials and other chemicals was used along with some simulated data. Assessing the use of residual variance functions when fitting non-linear toxicokinetic one-compartment models showed error margins depend on the actual model selected. Inspecting residual plots and error margins by means of PPD help to verify if a model fitted to the data was accurate. Adequate model selection may prevent either false positive or false negative conclusions regarding (nano)toxicity in various invertebrate organisms. It is essential that toxicokinetic models match the data and model assumptions should not be violated to ensure solid hypothesis testing.

## CRediT authorship contribution statement

**H.J. van Lingen:** Conceptualization, Data curation, Formal analysis, Investigation, Methodology, Software, Supervision, Validation, Visualization, Writing – original draft, Writing – review and editing. **E. Saccenti:** Conceptualization, Methodology, Supervision, Validation, Writing – review and editing. **M. Suarez-Diez:** Conceptualization, Funding acquisition, Project administration, Supervision, Validation, Writing – review and editing. **M. Baccaro:** Investigation, Methodology, Resources, Writing – review and editing. **N.W. van den Brink:** Conceptualization, Funding acquisition, Project administration, Supervision, Validation, Writing – review and editing.

## Acknowledgements

We thank Dr C.A.M. (Kees) van Gestel, emeritus Professor of Ecotoxicology of Soil Ecosystems at the Vrije Universiteit Amsterdam, for sharing various toxicokinetic datasets used for this study.

## Declaration of generative AI and AI-assisted technologies in the writing process

During the preparation of this work the author(s) used ChatGPT in order to improve the readability of the writing output generated by the authors. After using this tool, the authors reviewed and edited the content as needed and take full responsibility for the content of the published article.

