## Supplementary material for "Predicting uptake and elimination kinetics of chemicals in invertebrates: a technical note on residual variance modeling"

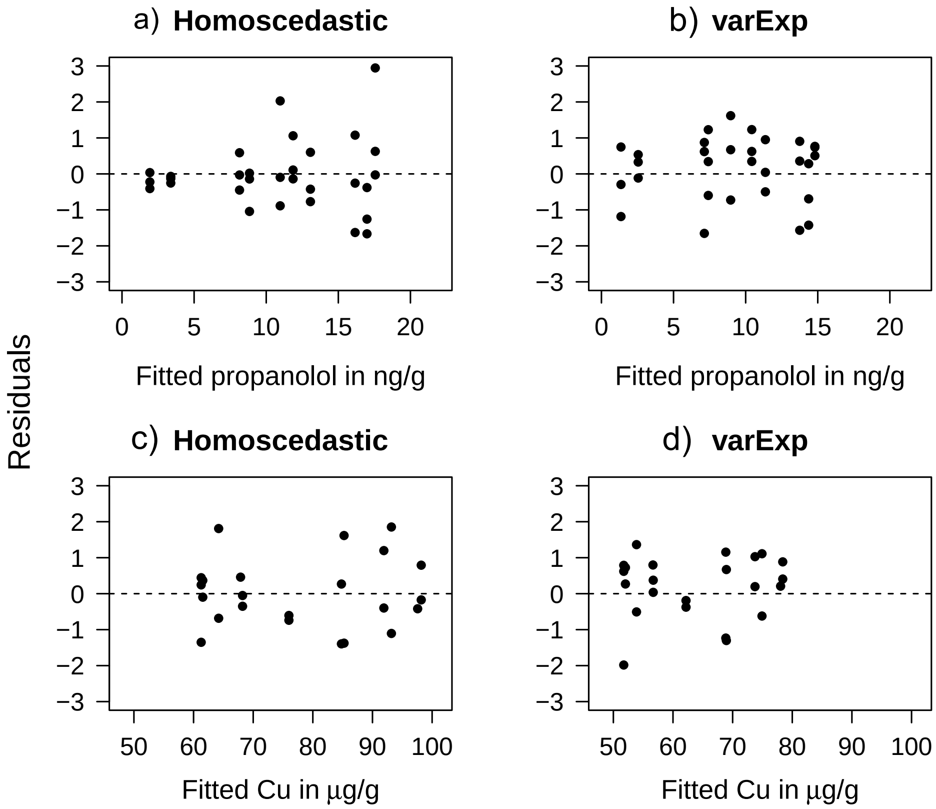


Figure A.5: Residuals of the fitted one-compartment toxicokinetic models fitted to the acquired data for *G. pulex* exposed to propanolol (panels a and b, respectively), and *F. candida* exposed to copper (panels c and d). Residuals are shown for the homoscedastic model fit (panels a and c) and the heteroscedastic model fits that used the exponential residual variance function (panels b and d).

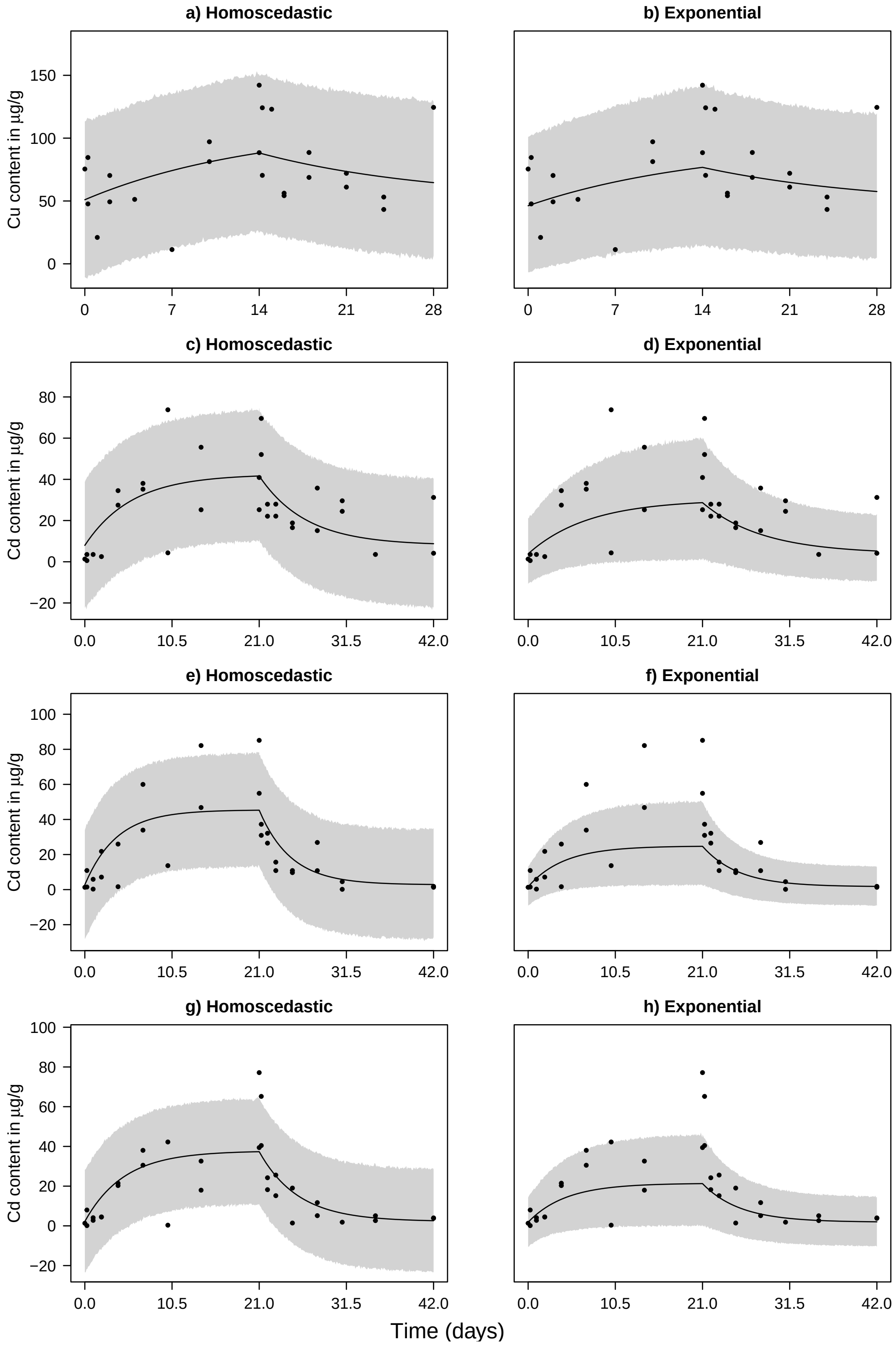


Figure A.6: Fitted one-compartment toxicokinetic models assuming homoscedastic residual variance (panels a and c) and heteroscedastic residual variance (panels b and d) using data acquired from (6; 27). Data points represent metal in contents in time in *F. candida* exposed to 25 mM Cu in soil with pH = 5.5 (panels a and b), to 20 mM Cd in soil with pH = 4.5 (panels c and d), pH = 5.5 (panels e and f) and pH = 6.5 (panels g and h). The black line and the grey shaded area indicate the mean and 95% credible intervals of the posterior predictive distribution, respectively. Estimated model parameters and fit statistics are shown in Suppl. Table A.2

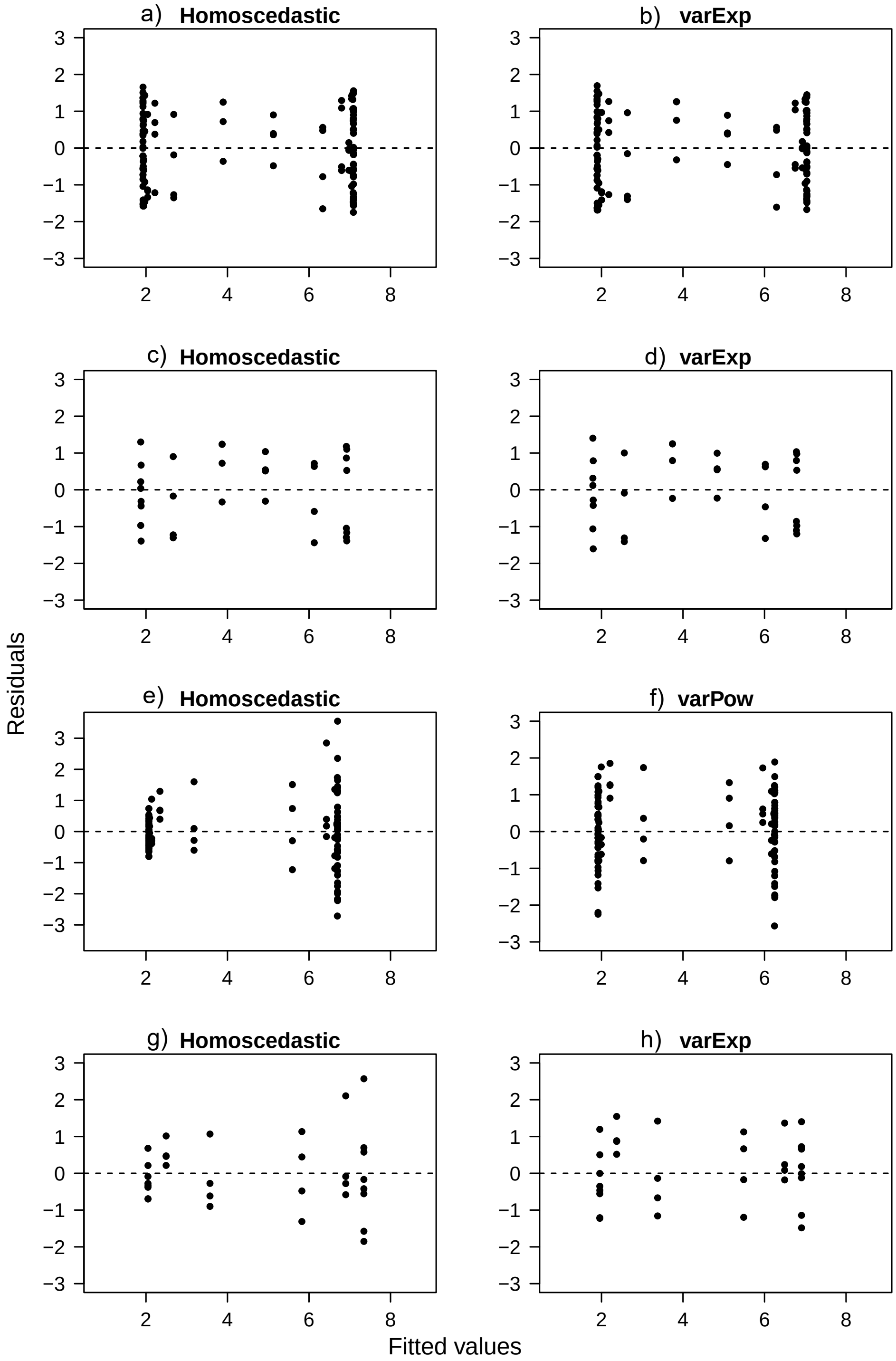


Figure A.7: Residuals of the fitted one-compartment toxicokinetic models to homoscedastically simulated data (panels a-d) and heteroscedastically simulated data (panels e-h). Residuals are shown for the homoscedastic model fit (panels a, c, e and g) and the heteroscedastic model fits that used the exponential residual variance function (panels b, d, f and h). Data points were retained for all 28 time units (full data; panels a, b, e and f) or only for time units 1, 2, 7, 14, 15, 16, 21 and 28 (reduced data; panels c, d, g and h).

Table A.2: Estimated parameters (mean ± standard deviation) for the fitted toxicokinetic homoscedastic and selected heteroscedastic models for predicting chemical contents along with log likelihood (logLik; quantifies the goodness of fit for which a higher score indicates better fit) and leave-one-out information criterion (LOOIC; quantifies the goodness of fit relative to the number of model parameters for which a lower score indicates better fit). $C_0$ represents the basal internal chemical contents in $\mu$g/g (mg or ng rather than $\mu$g may apply for certain systems), $k_1$ represents the uptake rate in g soil/g animal per time unit (e.g. hour, day) or L medium solution/g animal per time unit, $k_2$ represents the elimination rate per time unit, BAF represents the bioaccumulation factor in g/g or L/g. Model output is shown for datasets acquired from (6; 27)

| Model fit | $C_0$ | $k_1$ | $k_2$ | BAF | $\sigma_e$ or $\sigma_{\mathrm{e,uptake}}$ | $\sigma_{\mathrm{e,elimin}}$ or $\delta$ | logLik | LOOIC |
|---|---|---|---|---|---|---|---|---|
| *F. candida* exposed to Cu at 25 mmol/g of soil with pH = 5.5 | | | | | | | | |
| Homoscedastic | 51.0±8.2 | 0.15±0.06 | 0.072±0.040 | 4.2±26.5 | 30.1±4.6 | | -117 | 235 |
| Stratified | 51.4±8.3 | 0.15±0.06 | 0.074±0.041 | 4.2±36.9 | 31.5±7.3 | 29.4±6.2 | -119 | 237 |
| Fixed | 40.7±8.2 | 0.11±0.05 | 0.062±0.042 | 5.5±90.5 | 4.5±0.7 | | -123 | 246 |
| Exponential | 46.2±8.3 | 0.12±0.06 | 0.071±0.041 | 5.8±172.8 | 20.0±5.7 | 0.006±0.004 | -119 | 237 |
| Power | 47.8±8.5 | 0.14±0.05 | 0.069±0.041 | 5.8±81.6 | 17.1±7.3 | 0.17±0.12 | -119 | 237 |
| *F. candida* exposed to Cd at 20 mmol/g of soil with pH = 4.5 | | | | | | | | |
| Homoscedastic | 7.95±3.47 | 0.34±0.07 | 0.18±0.04 | 1.95±0.37 | 15.4±2.1 | | -126 | 253 |
| Stratified | 8.35±3.46 | 0.34±0.08 | 0.18±0.04 | 1.94±0.36 | 17.2±3.4 | 13.8±2.6 | -127 | 255 |
| Fixed | 1.85±1.27 | 0.22±0.06 | 0.14±0.04 | 1.66±0.34 | 3.4±0.5 | | -123 | 246 |
| Exponential | 3.84±2.43 | 0.20±0.07 | 0.14±0.05 | 1.50±0.36 | 6.3 ± 2.1 | 0.029±0.011 | -122 | 244 |
| Power | 1.99±2.04 | 0.20±0.08 | 0.13±0.04 | 1.52±0.46 | 3.5±2.6 | 0.57±0.25 | -124 | 247 |
| *F. candida* exposed to Cd at 20 mmol/g of soil with pH = 5.5 | | | | | | | | |
| Homoscedastic | 2.74±2.09 | 0.62±0.11 | 0.26±0.04 | 2.36±0.28 | 15.6±2.1 | | -127 | 254 |
| Stratified | 2.90±1.85 | 0.57±0.10 | 0.29±0.04 | 1.94±0.23 | 21.9±4.4 | 7.28±1.52 | -122 | 245 |
| Fixed | 0.60±0.54 | 0.26±0.07 | 0.25±0.04 | 1.04±0.25 | 4.0±0.6 | | -122 | 243 |
| Exponential | 1.65±1.20 | 0.30±0.09 | 0.23±0.05 | 1.28±0.27 | 5.0±1.3 | 0.033±0.009 | -115 | 230 |
| Power | 0.79±1.00 | 0.26±0.20 | 0.26±0.05 | 1.05±0.80 | 4.4±3.0 | 0.56±0.32 | -123 | 247 |
| *F. candida* exposed to Cd at 20 mmol/g of soil with pH = 6.5 | | | | | | | | |
| Homoscedastic | 2.20±1.67 | 0.44±0.07 | 0.21±0.03 | 2.09±0.25 | 12.9±1.7 | | -130 | 260 |
| Stratified | 2.15±1.63 | 0.47±0.08 | 0.23±0.03 | 2.10±0.24 | 15.4±2.9 | 10.1±2.0 | -132 | 264 |
| Fixed | 1.10±0.80 | 0.074±0.035 | 0.21±0.05 | 0.36±0.16 | 4.2±0.6 | | -131 | 261 |
| Exponential | 1.86±1.24 | 0.25±0.08 | 0.22±0.04 | 1.15±0.31 | 5.6±1.3 | 0.031±0.010 | -122 | 245 |
| Power | 0.89±1.31 | 0.15±0.19 | 0.21±0.05 | 0.68±0.86 | 4.5±4.6 | 0.66±0.43 | -133 | 266 |
